# Assessing group size and the demographic composition of unhabituated northern muriqui (*Brachyteles hypoxanthus*) using non-invasive biomonitoring

**DOI:** 10.1101/2024.12.13.628264

**Authors:** Mariane C. Kaizer, Naiara G. Sales, Thiago H. G. Alvim, Karen B. Strier, Fabiano R. de Melo, Jean P. Boubli, Robert J. Young, Allan D. McDevitt

## Abstract

Obtaining accurate population measures of endangered species is critical for effective conservation and management actions and to evaluate their success overtime. However, determining the population size and demographic composition of most canopy forest-dwelling species has proven to be challenging. Here, we apply two non-invasive biomonitoring methods, arboreal camera trap and genetic tagging of faecal samples, to estimate the population size of a critically endangered primate, the northern muriqui (*Brachyteles hypoxanthus*), in the Caparaó National Park, Brazil. When comparing population sizes between camera trapping and genetic tagging, the genetic tagging survey estimated fewer individuals for one of the muriqui groups studied but showed slightly higher population size estimates for the other group. In terms of the cost-efficiency of both methods, arboreal camera trapping had high initial costs but was more cost-effective in the long-term. Genetic tagging on the other hand did not require expensive equipment for data collection but had higher associated expenses for laboratory consumables and data processing. We recommend the use of both methods for northern muriqui monitoring and provide suggestions for improving the implementation of these non-invasive methods for future routine monitoring. Our findings also highlight the potential of arboreal camera trapping and genetic tagging to other arboreal mammals in tropical forests.

## 1 INTRODUCTION

Anthropogenic activities are leading to increasing threats to the world’s primates. It is estimated that 75% of all primate species have declining populations, and a major extinction event may soon occur if effective conservation strategies are not implemented (Estrada et al., 2017). To access the conservation status of a species, accurate information on population size(s) and trends over time are required (Mace & Lande, 1991; IUCN, 2017) to allow for effective management actions and an evaluation of their success over time (Nichols & Williams, 2006; Schipper & Rovero, 2017; Strier et al., 2017).

Monitoring species populations’ trends depends on reliable methods that can be replicated across different sites of species occurrence (Beaudrot et al., 2016; Steenweg et al., 2018). For most primate species, demographic trends are typically monitored by direct observation of habituated groups (Clutton-Brock & Sheldon, 2010; Williamson & Sheldon, 2010). However, this method is time-consuming and requires trained personnel, has high financial investment, and may have ethical issues (Fedigan, 2010; Williamson & Sheldon, 2010). Habituation may lead to disease transmission and increase the risks of poaching and is not recommended in places with pressure from hunters (Fedigan, 2010; Williamson & Sheldon, 2010; Strier et al., 2017). In addition, in remote forests or difficult-to-access areas, the implementation of this type of systematic monitoring may not be feasible. For canopy forest-dwelling species, this challenge is even more pronounced.

Recent technological advances in biomonitoring, such as camera trapping (McCarthey et al., 2018) and genetic tagging (Lamb et al., 2019), offer non-invasive alternative approaches to direct observation. Camera traps have been used to survey and monitor a wide range of taxa (Burton et al., 2015), including primates (Pebsworth & Lafleur, 2014). Standardization of field protocols has also allowed for the application of camera traps in long-term monitoring of a target species in different sites simultaneously (Scotson et al., 2017; Steenweg et al., 2017). Recently, camera trapping has been validated to survey and monitor chimpanzee communities, allowing for estimates of density through distance sampling (Cappele et al., 2019), spatially explicitly capture-recapture models (Després-Einspenner et al., 2017), and assessments of demographic composition and variation (McCarthey et al., 2018). Demographic and life history data have also been obtained for wild spider monkeys (*Ateles belzebuth*) through the monitoring of a geophagy site in the forest using camera traps (Galvis et al. 2014).

Non-invasive genetic monitoring also allows for population size estimation (Schwartz et al., 2007; Arandjelovic & Vigilant, 2018; Lamb et al., 2019). Non-invasive samples (e.g., hair, faeces) can be used in genetic approaches to identify distinct genotypes, providing information on the minimum number of individuals in a given area (Waits & Paetkau, 2005; Guschanski et al., 2009). Genetic “censusing” or “tagging” can also produce a detection history of individuals that can be combined with analytical methods such as spatial capture-recapture (Royle et al., 2017), allowing for an estimation of population abundance or density and their temporal and spatial variation (Whittington et al., 2005; McCarthy et al., 2015). Considering its efficiency for population assessments in a range of taxa (Kohn & Wayne, 1997; Taberlet et al., 1997; Woods et al., 1999; Arandjelovic et al., 2010), genetic tagging has the potential to be an effective tool for primate surveys. Despite this, however, it has only recently started to be applied in primatology, with most of the studies focusing on apes (Arandjelovic & Vigilant, 2018).

In this study, we evaluated the use of an integrative approach of non-invasive methods (arboreal camera trapping and genetic tagging) to obtain data on the size and group composition of one population of a canopy forest-dwelling primate: the northern muriqui (*Brachyteles hypoxanthus*), which is a critically endangered primate endemic to Brazil’s Atlantic Forest (de Melo et al., 2021). The need to obtain reliable and up-to-date demographic data on muriqui populations was highlighted in the National Action Plan for the Conservation of Muriquis (Jerusalinsky et al., 2011). Moreover, a more recent study has been published defining prioritized areas and intensity for systematic demographic monitoring of wild muriqui populations (Strier et al., 2017). As noted by Strier et al. (2017), in some of the areas it is not feasible or advisable to implement a systematic monitoring programme, and habituation may impose potential risks to the muriqui population. Therefore, non-invasive biomonitoring should be tested to improve species/population evaluations.

Our aim was to test the effectiveness of arboreal camera trapping and genetic tagging to assess the group size and demographic composition of unhabituated muriquis. The aims were threefold: 1) evaluate effectiveness of arboreal camera traps to collect demographic measures, including age-sex composition; 2) provide an assessment of group size based on arboreal camera trap data and genetic tagging; 3) compare the minimum group size of muriquis obtained from arboreal camera trapping with those obtained from genetic tagging, and the associated costs of each method. A better understanding of the aforementioned non-invasive methods to survey unhabituated muriquis in a remote rainforest will inform their potential and limitations for future monitoring programmes.

## 2 MATERIAL AND METHODS

### 2.1 Study site

This study was carried out at Caparaó National Park (PNC), located on the border of the states of Minas Gerais and Espírito Santo, in southeastern Brazil (20°37’ and 20°19’ S; 41°43’ and 41°55’ W). The 31,853 ha protected area was identified as one of the priority areas for the conservation of the northern muriqui (*B. hypoxanthus*) as it represents the highest altitudinal range of the species (Strier et al., 2017). The Park comprises a chain of mountains (Caparaó massif) that divides the area into two sides, east and west (Fig. 1), with altitudes ranging from 630 m to 2892 m above sea level (ICMBIO 2015). The landscape on the Caparaó massif includes rainforest in the lower altitudes and mainly altitude grasslands above 1900 m (Veloso et al., 1991). This study focused on two unhabituated groups of northern muriqui: one recently discovered group (VA: Aleixo Group) that inhabits a forest valley of ∼350 ha in the west area of the park (Kaizer et al. 2016), and another group that inhabits a valley of ∼1400 ha in the east area of the park (VSM: Santa Marta Group). During the study period, only one muriqui group inhabited the west valley, known as Aleixo valley, and no records of muriquis had been reported in adjacent valleys at the time of this research (Projeto Muriquis do Caparaó, 2020). For the east valley (Santa Marta valley), only one group was known, but other muriqui groups were found in immediately adjacent valleys. Due to the mountainous landscape, the muriqui group on the west side, VA group, did not have contact with the group on the east side, VSM group (Fig 1). While the east valley is characterized by continuous Montane Dense Ombrophilous forest, the west valley is characterized by Montane Seasonal Semideciduous forest (ICMBIO, 2015). Along the Caparaó massif, the climate type is “Cwc” (humid subtropical) according to the Köppen (1936) classification with a dry and cold season from May to September (Alvares et al., 2014). The average annual temperature and annual rainfall are 9.4^0^ C and 1,300 mm, respectively (Alvares et al., 2014).

**Figure 1.**
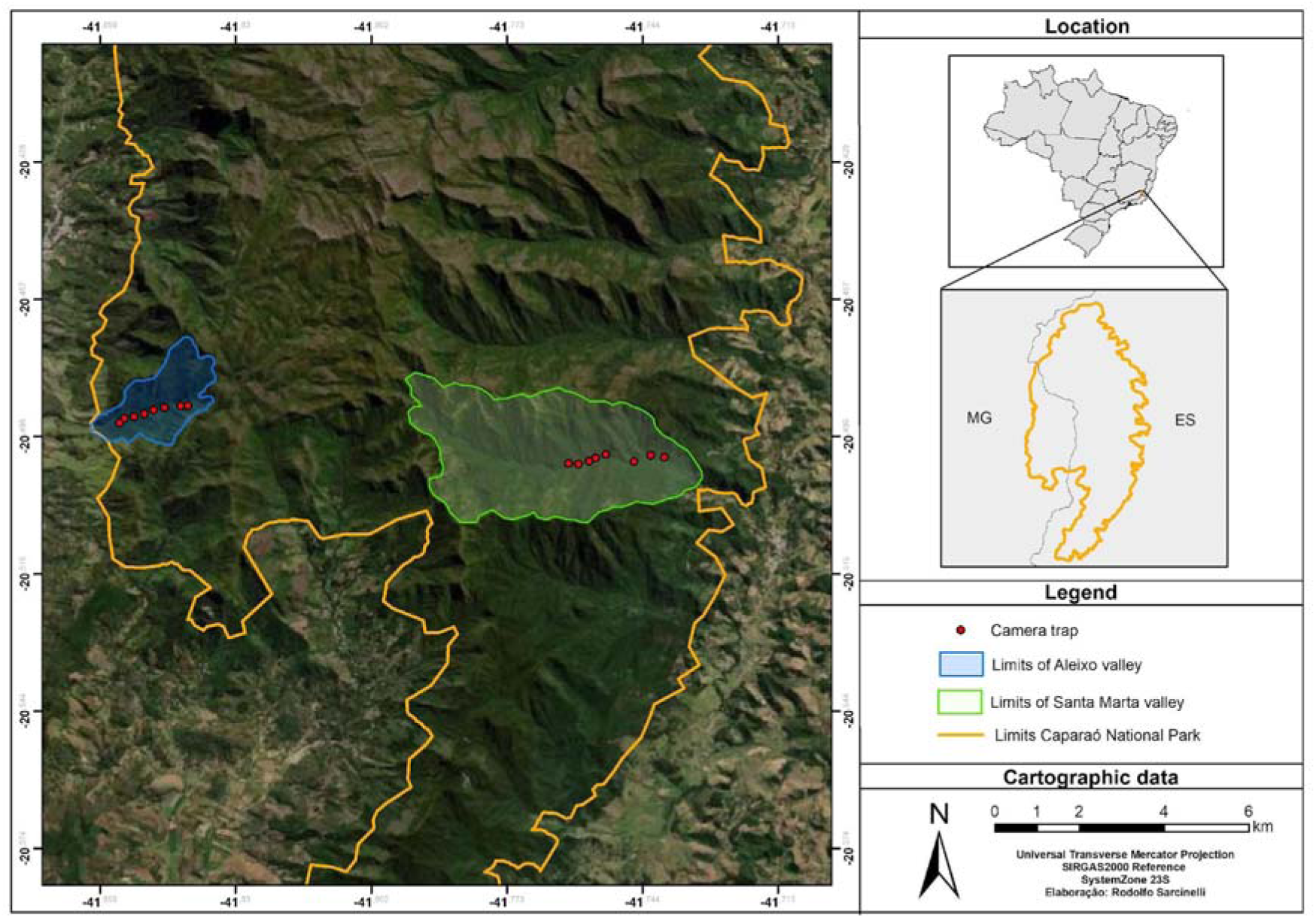
Map of Caparaó National Park (PNC), southeast Brazil, including locations of arboreal camera traps (red dots) in the two study sites: VA (left, blue) and VSM (right, green).

### 2.2 Arboreal camera trapping

Over a 12-month period, Jan-Dec 2017, eight infrared camera traps (Bushnell Trophy CamTM, #119774C; Bushnell Outdoor Products, USA) were deployed in the forest canopy (mean height: 11 m ± SD 2.5, range = 7.5-16 m) along a 2-km linear array spaced on average 255.7 m ± 66.9 (SD) in each site (Fig. 1). The remoteness of the areas hampered the use of a grid system; thus, a linear array was implemented. Each linear array was deployed across the forest valley, parallel to the valley’s main stream, potentially, crossing the territory range of muriquis. To ensure that groups were re-sampled in an unbiased manner (Arandjelovic et al., 2010), linear arrays covered a distance potentially no longer than the diameter of the home range of muriqui groups (122 – 406 ha, Lima et al., 2019), so that the same group was sampled on different occasions. Camera trap placement in the canopy followed protocol described by Kaizer et al. (2022; a more detailed description is included in Supplementary Material). Arboreal camera traps were set to work 24 hr/day in hybrid mode; that is, triggered to take two consecutive photographs (8MP) followed by one 30-s video (HD, 1280 x 720 pixels).

### 2.3 Camera trap data analysis

To investigate the group size and demographic composition of the unhabituated northern muriquis population we used units of temporal events obtained by the arboreal camera trap data. The adoption of units of temporal events from camera trap data has already been used by McCarthy et al. (2018) to define the party composition of habituated chimpanzees (*Pan troglodytes*). Since we set the camera traps to hybrid mode, which is characterized by a set of still pictures and a video that starts straight after, we defined a subevent as a set of two photographs and one video (following Gregory et al. (2014) for a set of 3 pictures). Thus, if an individual has been captured in a picture and in the subsequent video it was counted only once. The temporal event was defined as any subevents obtained at the same arboreal camera trap location occurring within 15 min of another on the same day (McCarthy et al., 2018). Therefore, all individuals passing in front of the same camera trap within the 15 min event have been counted.

To define independence between camera traps and to avoid overestimating the minimum size of muriqui groups, when muriquis were recorded in more than one camera trap in the same day, we selected for analysis that camera which recorded the largest group size (McCarthy et al., 2018). Whenever the records were obtained within an interval of 10 min of difference between one camera, we considered them as independent events.

### 2.4 Identification of group size and composition in arboreal camera trap data

To define the demographic composition of the population based on canopy camera traps, we counted the number of individuals classified in different ages and/or sex classes. Muriquis can be easily distinguished by their sex and age class due to the physical features that correspond to their developmental stages (Fig. 2). Adults can reach as much as 15 kg (Aguirre, 1971), and their sex can be easily differentiated due the size and shape of their genitalia (Strier, 1987). Even for juveniles, sex can usually be determined based on genitalia shape and positioning (Strier et al., 2006) and physical features associated with behavioural development described for muriquis may allow for the identification of infants (Odália-Rimoli, 1998).

**Figure 2.**
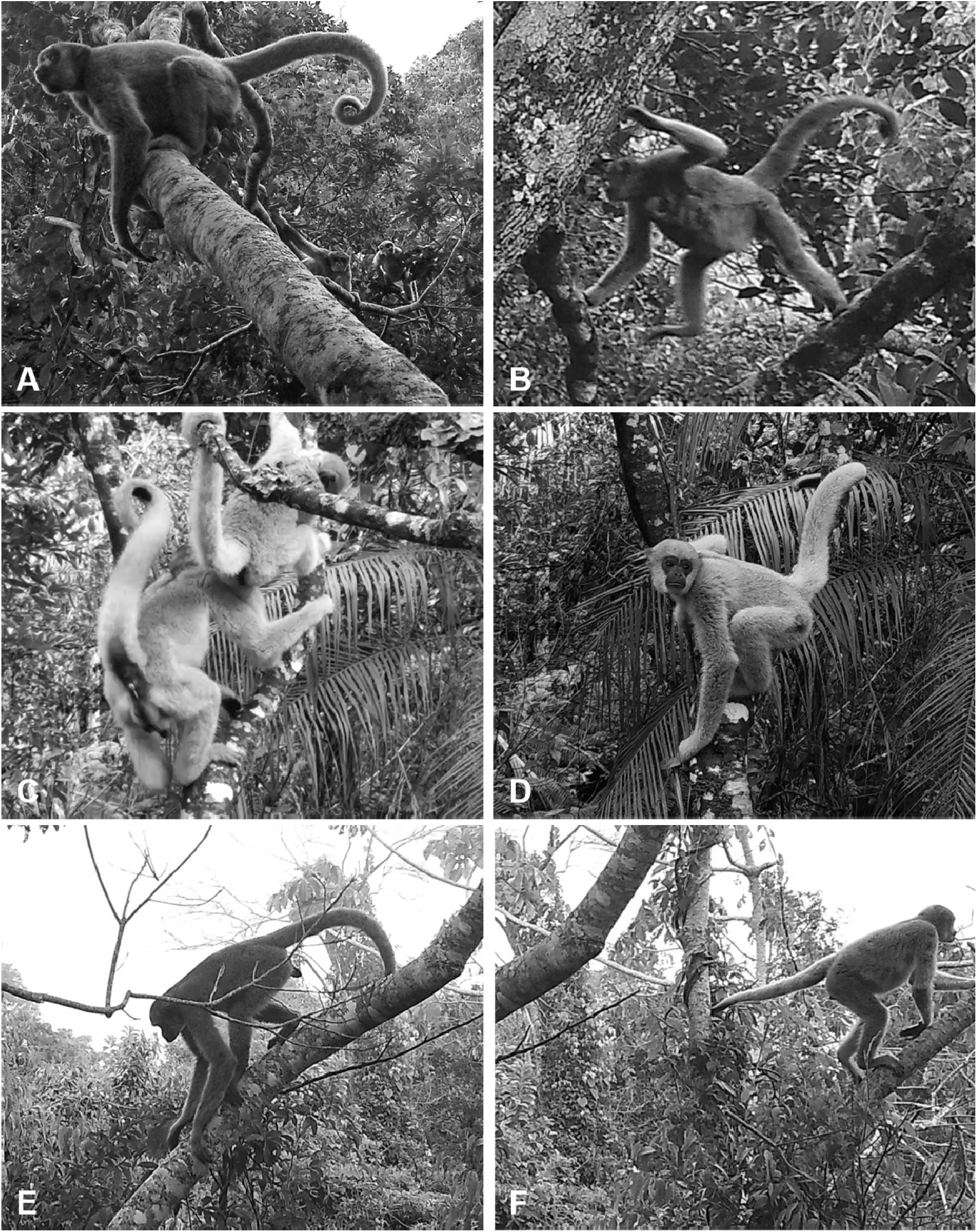
Northern muriqui (*Brachyteles hypoxanthus*) recorded in canopy camera traps at Caparaó National Park, Brazil. From top to bottom: (A): two adult males on the left and one subadult in the back of the picture, (B) adult female carrying a new-born ventrally, (C) adult and juvenile males, (D) juvenile male, (E) adult female, (F) juvenile female.

Camera trap data were analysed by an experienced northern muriqui researcher (MCK), following the protocols established by Strier et al. (2017, S1 Table) to define the age-sex class of individuals. Based on the quality of the camera trap data, associated physical features and behaviour of muriquis of a specific age class (e.g., infants are carried ventrally or dorsally by their mothers which prevents their sex identification), we extracted the number of individuals in each demographic class as follows: immatures dependent: 0-2 years; immatures independent: 2-5 years; subadults (F/M): ∼5-8 years; adults (F/M): <8years (for more details, see S1 Table and Fig. A-R in S1 File in Strier et al., 2017). The number of individuals whose age-sex class could not be defined with precision due to the quality of images/video or due to the position of the animal was classified as “unidentified”.

Following McCarthy et al. (2018), we inferred the population demographic composition of unhabituated muriquis as follows: i) by examining the size and composition of the largest subgroup passing in a single camera trap event (i.e. within 15 min); and ii) by calculating the largest number of individuals in a given age-sex class obtained in an event (e.g. the largest number of adult females detected together), then combining the maximum number of distinct individuals in each category as a proxy measure for total group composition.

### 2.5 Genetic sampling

We conducted a repeated systematic survey for non-invasive faecal samples, during one week per month, in the two sites during the same camera trap survey period (over Jan-2017 to Jun-2017), alongside a pre-existing trail used as a transect. Additional samples were obtained, opportunistically during other fieldwork activities, between January and February of 2018. Date, time, and geographic coordinates were noted for each sample collected (Table A1).

Faecal samples were collected fresh and immediately stored in RNALater (Ambion), using a 1:1 ratio (sample: nucleic acid stabilization buffer). A detailed description of all laboratory protocols (DNA extraction, mitochondrial and microsatellite amplification) and bioinformatics is included in the Supplementary Material. In brief, after DNA extraction the mitochondrial control region haplotypes were obtained, and a molecular species identification was conducted through a BLAST search in the NCBI database. Then, sequences were aligned using the CLUSTAL W and visually inspected in MEGA version X.

For the genotyping analyses using the nuclear microsatellites, a panel of 14 loci was screened through multiplexed reactions (Table A2; Sales et al. 2024). All analysed samples were genotyped at least three times for all loci, to minimize genotyping errors. Automated allele calling was conducted in GeneMapper v.4.1 (Applied Biosystems) and further visually checked for genotype determination. Genotypes were independently determined twice for all samples per replicate (Stojak et al., 2016). Homozygote genotypes were considered when scored on all replicates or in at least two out of the three replicates when one replicate amplification failed. Heterozygote was accepted if scored on two out of three replicates, if none of these criteria was met, alleles were classified as missing data.

### 2.6 Group size data from genetic sampling

To estimate group size, we used the number of individuals identified by genetic analyses based on mitochondrial DNA (control region) and genotypes obtained for a panel of 14 microsatellite loci. The probability of identifying (PI) measures for unrelated (PI*_ave_*) and siblings (PI*_sibs_*) were calculated in GenAlex (Peakall & Smouse, 2006). Genetic tagging of individuals was based on the search for matching genotypes. Distinct individuals were considered when either showing a different mtDNA haplotype and same/distinct microsatellite genotypes; or having the same mtDNA haplotype and distinct microsatellite genotypes. Duplicates (i.e., samples obtained from the same individual) were considered in the presence of a match for both microsatellite genotypes and mtDNA haplotypes. All samples identified with the same haplotype and a matching score ≥87.5% loci were re-examined for possible genotyping errors.

Since unhabituated muriquis were sampled and the genetic sampling was conducted in distinct sampling sessions, individuals could be sampled or “recaptured” several times. We considered the independence of genetic sampling events as follows (Arandjelovic et al., 2010; McCarthy et al., 2015): i) distinct samples from the same individual were considered independent, if they were separated geographically and/or temporally on distinct days; ii) distinct samples from the same individual were considered duplicate, if they were collected on the same day and at a distance apart of <3479 m apart, based on the maximum travel distance reported for habituated muriquis within one day (Lima et al., 2019).

### 2.7 Survey methods costs

We calculated total purchase costs, labour expenses, data processing and on-site logistical costs associated with canopy camera trap survey and genetic tagging survey methods for our study. To increase the time- and cost-efficiency of fieldwork, the collection of faecal samples for genetic tagging was carried out in parallel to other fieldwork activities, but for comparison purposes, here we assumed independence between survey methods and calculated costs separately. Labour associated with the lead author’s time performing camera trapping and genetic analyses were not computed, thus we provided only costs required for data processing without including data analysis. We calculated costs for canopy camera trapping and for genetic monitoring using UK prices in 2018.

## 3 RESULTS

Across an effort of 2,613 canopy camera trap days, we obtained 948 records (pictures and videos) of northern muriquis. These records resulted in a total of 148 temporal events (15 min, Fig. 3) of which 95 corresponded to the VA site, and 53 corresponded to the VSM site. An average of 2.45 ± 2.26 (SD, range: 1 – 10) subevents (2 pictures and 1 video) per event was obtained in the VA site and an average of 1.70 ± 0.99 (SD, range: 1 – 5) in VSM.

**Figure 3.**
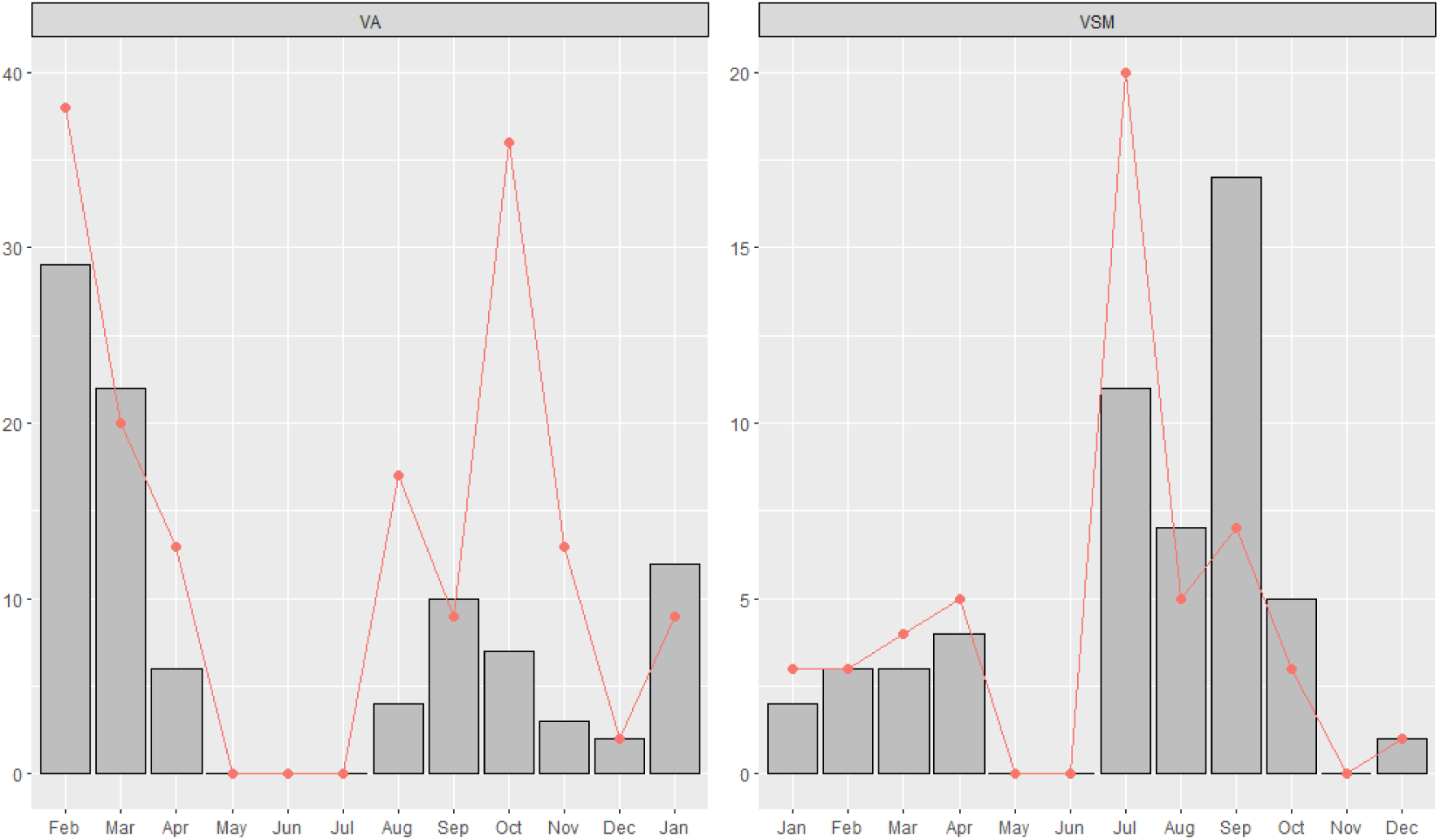
Seasonal variation of number of temporal events (15 min; grey bars) and maximum group-size of northern muriqui (*B. hypoxanthus*) recorded in the same arboreal camera trap (pink dots), in two distinct sites within Caparaó National Park (PNC), Brazil. Surveyed over a 12-month period (VA: Feb-2017 to Jan-2018; VSM: Jan-2017 to Dec-2017).

Camera traps detected muriquis on 80 distinct days across the survey period, from which nine days corresponded to simultaneous records of the species in the two sites. Muriquis were recorded across the 2-km array of eight arboreal camera traps in the VA site, but only in five camera trap sampling points in the VSM site. In the VA site, muriquis were recorded in canopy camera traps for 52 days of which 13 days corresponded to multiple records of muriquis in more than one camera (average 2.3 ± 0.5 SD, range: 2 – 3). This resulted in a range of 606.2 m ± 352 (SD, range: 255 – 1270 m). By contrast, in the VSM, muriquis were recorded in more than one camera (2 cameras) on only two of the 37 days of records, over a range of 255 m.

### 3.1 Group size and demographic composition

The largest subgroup sizes captured on an arboreal camera trap in a single temporal event (15 min), comprised 36 muriquis in October 2017 at the VA site and seven individuals in July 2017 at the VSM site. However, when analysed the total number of observed muriquis passing in the same camera trap in subsequent temporal events, this number increased to 38 muriquis at the VA site and to 20 at the VSM site (Fig. 3). Across all events, the summed maximum numbers of individuals seen in each demographic class were 40 distinct muriquis in the VA site, and 23 muriquis in the VSM (Fig. 4). The maximum number of mature individuals was 21 on the VA site (Male: 12, Female: 6, Unidentified: 3), and 16 on the VSM site (Male: 11, Female: 2, Unidentified: 3; Fig. 5).

**Figure 4.**
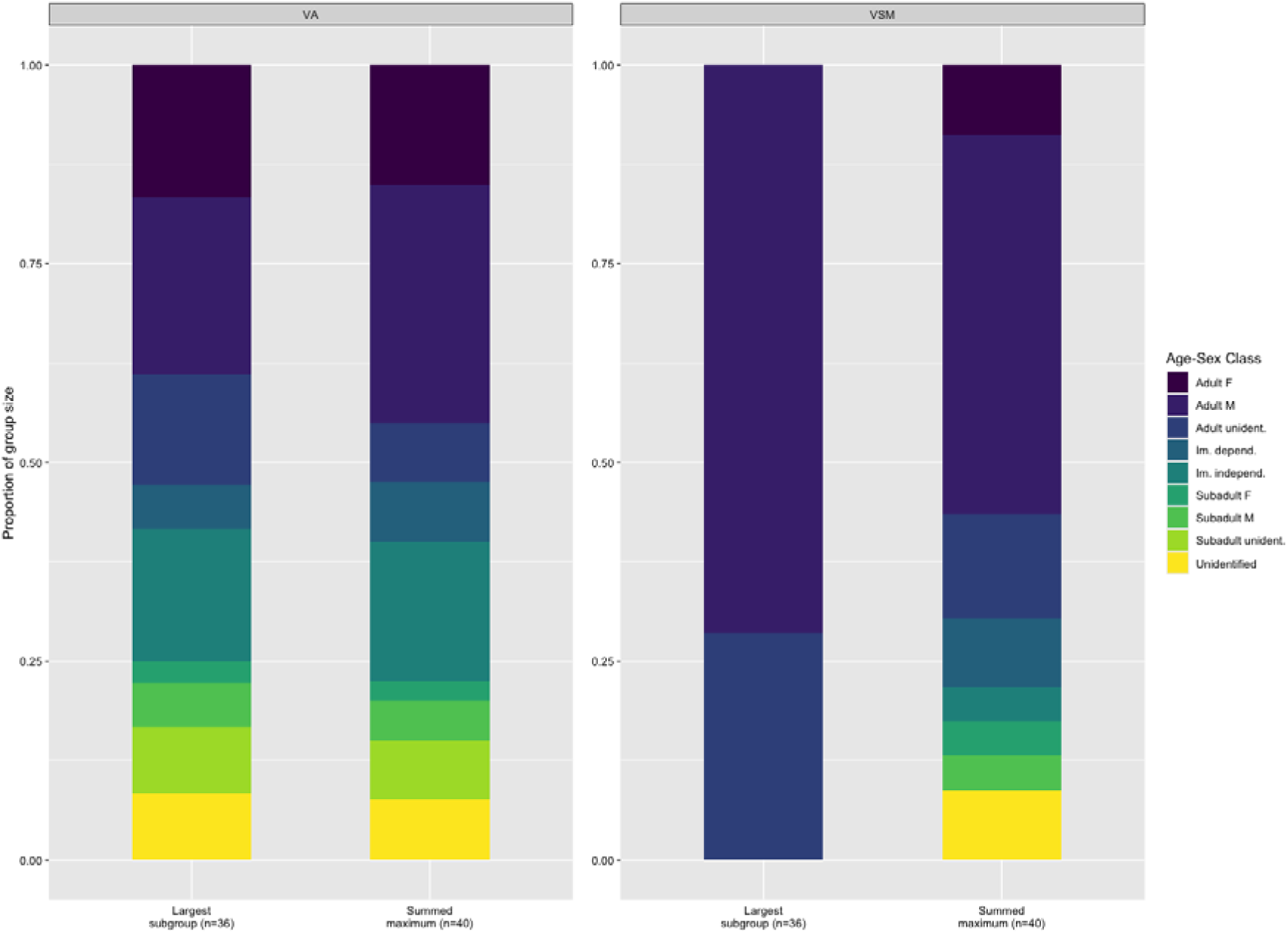
Demographic composition of two northern muriqui (*B. hypoxanthus*) social groups at Caparaó National Park, Brazil, inferred by proxy estimates of the largest subgroup size recorded by a single arboreal camera trap event and by the summed maximum numbers of muriquis in each demographic class (Im. dependent: immature dependent; Im. Independ.: Immature independent; M: male, F: female, Unident: unidentifiable). VA group on the left and VSM on the right.

**Figure 5.**
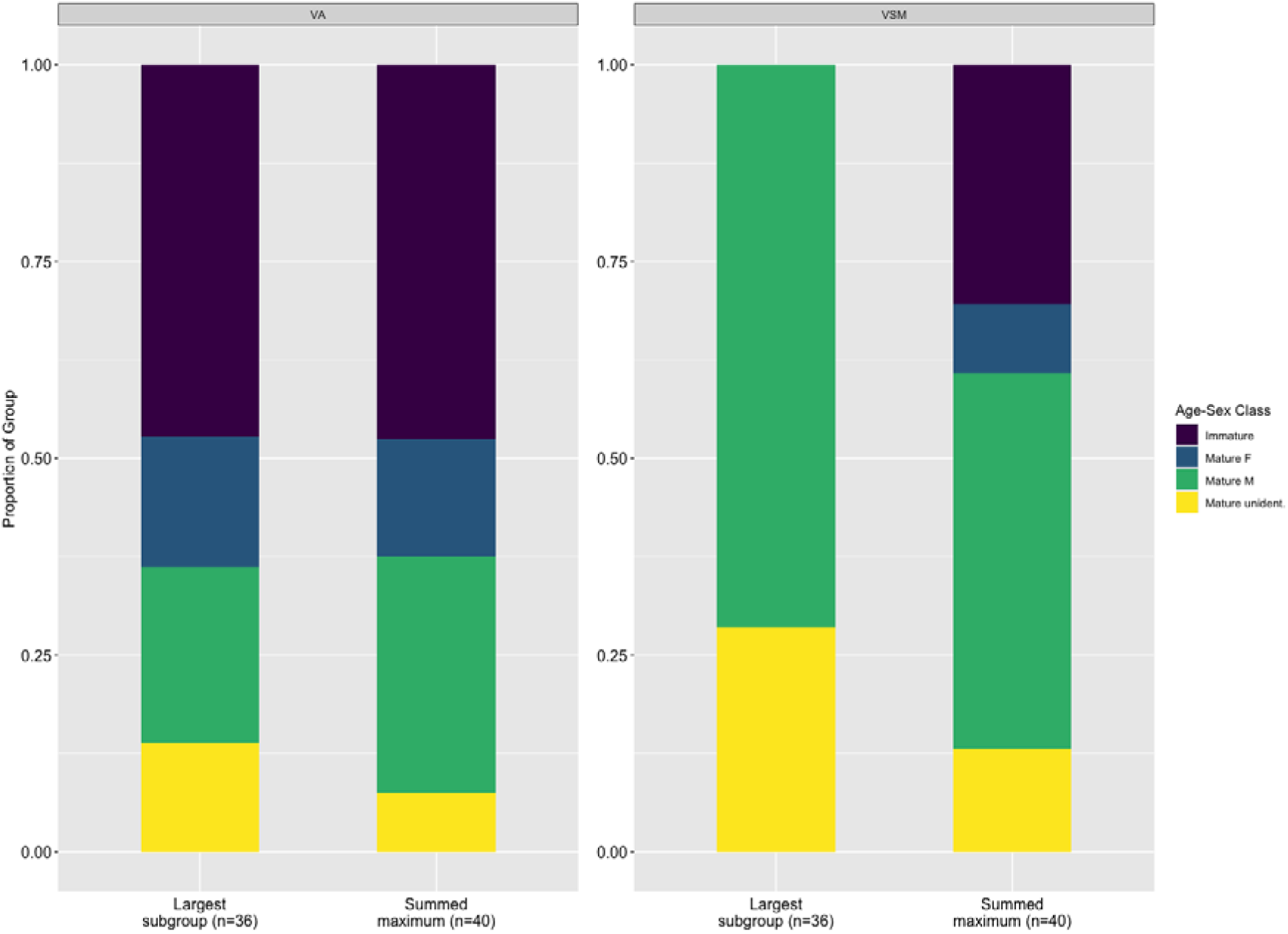
Proportion of mature and immature individuals in the two northern muriqui (*B. hypoxanthus*) social groups at Caparaó National Park, Brazil. Measures inferred by proxy of the largest subgroup size recorded by a single arboreal camera trap event and by the summed maximum numbers of muriquis in each demographic class (M: male, F: female, Unident: unidentifiable). VA group on the left and VSM on the right.

At least three adult females were carrying dependent immature individuals in the VA site, and two in the VSM site. Of these records, one represented a newborn infant recorded on the VSM site, indicating that at least one birth occurred there during the study period. The adult female was recorded carrying the infant ventrally and nursing it during July 2017.

### 3.2 Genetic tagging

We collected a total of 60 faecal samples across the period of the survey. Different shapes and small sizes of some faecal samples suggested individuals of different ages were sampled. An average of 3.83 ± 4.13 SD (range: 1 – 15) faecal samples were collected per day over 12 days at the VA site. In the VSM site, an average of 2.14 ± 1.68 SD (range: 1 – 5) faecal samples were collected in seven days. Putative muriqui samples were genotyped using the 14 sets of microsatellite loci and misclassified samples (n=3) were excluded. PI*_ave_* across the 14 loci was <0.001 for the VA and VSM groups, and PI*_sibs_* was 0.000030 and 0.000035 for VA and VSM groups, respectively. Therefore, this panel of 14 microsatellite loci had sufficient power to distinguish between individuals in each of these populations. The 46 samples collected from the VA group yielded 44 usable genotypes representing 42 genetically distinct individuals. The 14 faecal samples collected from the VSM group yielded 13 usable genotypes and resulted in 13 unique individuals.

The genotypes of the 55 resulting individuals were on average 83.4% complete (average of 11 loci genotyped per sample). Since a few samples presented the same mtDNA haplotype and were differentiated at a minimum of one locus (which were rechecked after genotyping matching analysis), we are confident that we did not incorrectly count the number of individuals.

Two of the 57 usable genotypes represented duplicate samples (VA=1, VSM=1), which were sampled on the same day separately <3479m. Non recapture events were noted.

### 3.3 Canopy Camera Trapping *vs* Genetic Tagging Surveys

Group size estimates for the VA group were relatively similar among canopy camera trapping (CCT) and genetic tagging (GT) survey methods (CCT: 40 individuals; GT: 42 individuals). But the genetic tagging survey provided a lower estimate of group size than that of canopy camera trapping for the VSM group (CCT: 23 individuals; GT: 13 individuals).

Table 1 shows the financial costs incurred in conducting the canopy camera trapping survey and the genetic tagging survey for group size estimation of northern muriqui in this study. The 12-month canopy camera trapping survey costs (£9,774.40) were a little lower than the costs for the genetic tagging survey (£11,402.68). Our results showed that the initial equipment costs (mainly climbing equipment and cameras) for arboreal camera trapping were counterbalanced by the reduced effort in field labour during data collection. The costs associated with the genetic tagging survey comprise mainly laboratory consumables and DNA processing, which are associated with the number and quality of samples. Maintaining a sample collection team in the field also added substantial costs to the genetic survey, which required more visits per site.

**Table 1.** Cost comparison between arboreal camera trapping survey and genetic tagging survey for group size estimation of northern muriquis (*Brachyteles hypoxanthus*) in the Atlantic Forest of Brazil. Prices (UK) are based on 2018 year.

|  | Descriptor | Survey Cost (GBP) |
| --- | --- | --- |
| **Canopy Camera Trapping** |  |  |
| Equipment | Climbing Equipment | £1517 |
|  | Cameras (16 x Bushnell Trophy Cam) | £2756 |
|  | Memory cards 32GB | £260 |
|  | GPS Garmin (+ battery charger) | £220 |
| Training costs | BCAP<sup>1</sup> Climbing course | £595 |
| Vehicle | Rental | £945 |
|  | Fuel | £300 |
| Consumables | Batteries | £289 |
| Data storage | External hard drive (5 TB) | £120 |
| Field labour | Set up (8-cameras array/site) | £512 |
|  | Review – 4 visits | £1280 |
| Field station costs | Accommodation and food | £240 |
| Camera trap data processing | Images and videos | £740 |
| **Overall costs of survey** | **12-month survey** | **£9,774** |
| **Genetic Tagging** |  |  |
|  | GPS Garmin (+ battery charger) | £220 |
|  | Faecal sample preservation (RNAlater) | £365 |
|  | Field consumables (plastic tubes, gloves) | £50 |
| Vehicle | Rental | £1323 |
|  | Fuel | £420 |
| Field labour | Faecal sample collection – 7 visits | £2240 |
| Field station costs | Accommodation and food | £280 |
| DNA processing | Laboratory consumables (tips, gloves, tubes, plates, reagents, DNA kit extraction) | £3000 |
|  | Primers | £431 |
|  | mtDNA sequencing | £693 |
|  | Microsatellite genotyping<sup>3</sup> | £2380 |
| **Overall costs of survey** |  | **£11,402** |

<sup>1</sup>BCAP: Basic Canopy Access Proficiency
<sup>2</sup>DNA processing assumes 80 samples
<sup>3</sup>Assuming 16 microsatellite loci

## 4 DISCUSSION

Non-invasive monitoring such as arboreal camera trapping and genetic tagging for population assessments of arboreal primates remain underused, despite their effectiveness having been demonstrated over the last 20 years (Arandjelojic & Vigilant, 2018; Kohn et al., 1999; Lamb et al., 2019; O’Connell et al. 2011). In this study, we provided evidence that both arboreal camera trapping and genetic tagging are promising tools for unhabituated arboreal primates, especially for populations of endangered species where the use of systematic demographic surveys may not be feasible.

This study focused on an unhabituated muriqui population, with an unknown population size, and therefore inferences regarding the similarity between the true number of individuals and both camera trap data and genetic tagging could not be made. Group size estimates varied between survey methods in one of the two muriqui groups sampled. Specifically, although estimates were relatively similar for the VA group, the group size estimation for the VSM group provided by the genetic tagging survey represented only 56.5% of that provided by the canopy camera trapping. This conservative estimate from genetic tagging may be a result of the difficulty of surveying immature muriquis, despite collecting faecal boluses of small sizes. Usually, genetic censuses of primates tend to underestimate younger individuals due to the difficulties in finding small size faecal samples in the field (McCarthy et al., 2015).

In comparison between the two sites, we found the largest group size in the VA site and the largest number of individuals captured on an arboreal camera trap in a single temporal event (15 min). The maximum number of faecal samples collected on a single day also suggested a similar pattern (VA: 1 – 15; VSM: 1 - 5). It is known that flexibility in group dynamics in muriquis is common (fission-fusion and cohesive groups) and might be a reflection of the differences in demography and ecological features between the sites (Strier & Mendes, 2012, 2016; Silva Junior et al., 2009; Strier, 2017). Although it is outside of the scope of our objectives here, these findings suggest that arboreal camera trapping may hold promise in detecting variations in group size and their associations with habitat structure and other ecological factors (McCarthy et al., 2018).

There was an absence of muriqui detections during the dry season months of May and June in both sites. Muriquis are a folivore-frugivore species known to use temporarily available resources (Strier, 1991), which might explain the absence of detections during these months. The tree *Mabea fistulifera* (Euphorbiacea), for example, is an important seasonal resource for muriquis in other populations (Mourthé et al., 2008). This pioneer species, which presents peak inflorescences in late April-May, is commonly found on the edge of forests that suffer severe disturbances or dominates the hilltops that suffer forest fires (Boubli et al., 2011). Therefore, the future use of arboreal camera trapping deployed in a grid or randomly distributed within the study site may contribute to our understanding of habitat use and resource distribution in remote areas.

The IUCN Standards and Petitions Subcommittee defines population size and its trends over time employing the number of mature individuals and its variation in a population/species (IUCN, 2017). Our findings revealed that both arboreal camera trapping and genetic tagging are potential tools to estimate the minimum population size of unhabituated muriquis. Furthermore, although arboreal camera trap data required some time investment to confidently classify the demographic class of individuals, the identification of mature individuals was relatively easy and did not require individual identity recognition (but artificial intelligence techniques could automate these processes in the future; Paulet et al., 2024). Arboreal camera trapping revealed a difference in the number of mature males and females in both muriqui groups, and allowed for the identification of immature dependents, providing inference of the number of reproductive females in the population. Additionally, despite the assumption of demographic closure required by estimating population methods (capture-recapture, Karanth & Nichols, 1998; Lukacs & Burnham, 2005) an adult female carrying a newborn infant was registered indicating the occurrence of birth events in the population. Furthermore, a recent study assessing the efficiency of camera traps for surveying demographic composition and variation in chimpanzees suggested the feasibility of the method to survey unhabituated populations (McCarthy et al., 2018). By matching data on party size and demographic composition among camera trap data and direct observation of habituated chimpanzees, camera trap data tended to underestimate party size but demonstrated proxy patterns of demographic composition (age/sex proportions) in the absence of individual identification (McCarthy et al., 2018).

The genotyping success (83.4%) based on faecal samples reported in our study was consistent with genotyping success reported for other unhabituated primate populations (chimpanzees: 77-83%, Moore & Vigilant et al., 2014; McCarthy et al., 2015). Despite the influence of a wide range of factors (e.g. time exposed to the environment, temperature, UV, microbial activity; Waits & Paetkau, 2005; Beja-Pereira et al., 2009), the genotyping success rate found in our study demonstrated the potential application of genetic tagging to assess canopy forest-dwelling primate populations in remote areas. Our results revealed a conservative estimate (minimum group size) of 42 individuals in VA and 13 individuals in VSM and identified two duplicate samples. Dating and locations of the faecal samples were key components to identify and understand duplicates. The two duplicate samples were collected on the same day within a very short distance; probably, because of the arboreal behaviour of muriquis (i.e. faecal samples can be fragmented into distinct pieces when hitting the ground). However, our sampling was conducted following a linear trail limiting the number and distribution of samples obtained; thus, preventing further detailed analyses such as mark-recapture analysis, due to the lack of ‘recaptures’. We suggest for future studies that defined grids should be considered where this is feasible, since muriquis usually present large home range sizes (406 ha on average, Lima et al., 2019) and move heterogeneously throughout the forest. A grid-based design with consistence effort may increase the sampling efficiency, minimizing the time effort and reducing costs (Arandjelovic & Vigilant, 2018). Additionally, the precision and accuracy of population size estimation increase with the number of repeated samples (Arandjelovic & Vigilant, 2018). Thus, 3-4 times the number of samples compared to the expected/potential number of individuals in the study population should be collected (Petit & Valiere, 2006; Arandjelovic, 2010; McCarthey et al., 2015). As a good starting point, a genetic monitoring programme could be implemented in a muriqui population of known size to determine the optimal sampling strategy and to validate capture-recapture methods to infer a better rule-of-thumb number.

Despite the limitations of our study design and the challenge to compare survey techniques when the true group size is not precisely known, our results demonstrated that genetic samples collected over 12 days resulted in a relatively close group size estimation to that of 12 months of arboreal camera trapping, at least for one of the groups. This finding strengthens our suggestion that a systematic genetic sampling study may be also a cost-effective tool for population size assessments of unhabituated muriquis, especially in rapid assessments (Hedges et al., 2013). Several studies have reported the efficiency of genetic tagging over traditional direct monitoring (McCarthey et al., 2015; Granjon et al., 2017) and other methods (Zhan et al., 2006; Hádjkova et al., 2009; Arandjelovic et al., 2010), suggesting that genetic counts may provide more precise and accurate counts (Solber et al., 2006). Moreover, genetic methods can also be used for sex identification (Di Fiore, 2005; Waits & Paetkau, 2005) and age class (Carroll et al., 2018; Cattet et al., 2018), and non- or minimally-invasively collected samples (e.g. hair) can be used to infer the reproductive status of individuals through hormones (Cattet et al., 2017). However, these analyses have not been applied in this study due to time and financial constraints.

Our cost comparisons show that despite the high upfront costs of equipment purchasing to implement the canopy camera trapping survey, the subsequent sampling efforts by this method are substantially less expensive. Therefore, indicating a better cost-effective survey, especially for long-term monitoring programmes (Burgar et al., 2018). Although non-invasive genetic tagging does not require expensive equipment for data collection, the costs of laboratory consumables and genetic data processing are high and will vary according to the number of samples and/or markers. Thus, studies that require repeated sampling sessions to collect a very large number of samples may become cost-prohibitive (Hedges et al., 2013). However, genetic tagging surveys may have reduced costs when combined with other field-based methods by sharing logistical efforts and costs (Guschanski et al., 2009). Good quality faecal samples may also decrease the costs of genetic analysis by minimizing genotyping errors and improving data quality (i.e., fewer repeat analyses; Beja-Pereira et al., 2009).

Here we have shown for the first-time an estimate of the minimum population size and demographic composition of two unhabituated muriquis groups. Our findings suggest the feasibility of arboreal camera trapping and, to a lesser extent, genetic tagging methods to fill the critical information gaps in extant muriqui populations, especially in those areas of geographic importance where systematic demographic monitoring is undesirable or poses risks to the population (Strier et al., 2017). Although both methods have intrinsic limitations (e.g., initial cost investment, time-consuming data analysis), the combined application of these non-invasive approaches may offer a range of opportunities to improve wild arboreal primate population monitoring. Finally, we observed that the total number of individuals differed greatly between sites, with the largest area presenting the smaller group size, and that the number of mature individuals differed between males and females in both groups. These empirical results raise concerns regarding the future reproductive potential of this population and highlight the importance of expanding our surveys to other sites within the park to provide a more realistic estimate of the entire muriqui population, and to inform conservation management strategies in this protected area.

## Supplementary material

Appendix A. Supplementary data

## Acknowledgements

We gratefully acknowledge the Parque Nacional do Caparaó/ICMBio, extending to its managers, staff, and surrounding community, for administrative and logistic support. We thank CLN, FGH, and RS for the valuable fieldwork assistance, DSF and AC for the project’s logistical support, and the MEG team, especially DS and SB, for their helpful support on laboratory and genetic analyses. The present study was carried out with all required permits (SISBio N. 54795, DEFRA: ITIMP17.1302, SisGen: A5C63DC) and ethical approval (STR1718-14).

## Declaration of competing interest

The authors declare no conflicts of interest.

## Funding

This work was supported by the Conselho de Aperfeiçoamento de Pessoal de Nível Superior, Brazil [CAPES, proc. BEX 1 298/2015-01] through the PhD scholarship to MCK; the Conservation Leadership Programme [grant N. 12455]; the MBZ Fund [grant N. 162512917]; the Idea Wild; and the Conquista Montanhismo.

## Notes

### Competing Interest Statement

The authors have declared no competing interest.

