## Supplementary Material for "Assessing group size and the demographic composition of unhabituated northern muriqui (*Brachyteles hypoxanthus*) using non-invasive biomonitoring"

**Material and Methods**

***Arboreal camera trap placement***

To install the camera trap in the forest canopy, we used rapid ascent/descent (RAD) and single rope (SRT) techniques (Maher, 2006) to climb the trees, and positioned the cameras to monitor potential arboreal pathways to detect animals in a neighbouring tree and/or along a horizontal branch in the same tree. Since the survey was conducted in a densely vegetated and continuous forest, the trees were chosen according to their safety to climb and the presence of connection between trees by the proximity of its branches forming an aerial pathway through the canopy (i.e. tree connectivity). Trees were connected to at least three other neighbour trees allowing the detection of potential arboreal mammals independently of species or animal’s age. No baits or lures were used to attract animals, and no fruiting trees were chosen as places for cameras, to avoid bias data. We placed the camera traps on or close to the tree’s trunk (to prevent movement due to the wind and to not interrupt arboreal pathways), without the use of brackets or mounts to secure the cameras (to prevent injuries to the trees and reduce costs). To prevent biases and to reduce exposition to sunlight, in each site we deployed all cameras faced to the same cardinal direction (VA = northerly direction, VSM = southerly).

Each camera was equipped with 32 GB memory card and eight AA alkaline batteries. Over a 12-month period of survey, we visited arboreal sampling points every three months to perform camera maintenance and exchange memory cards and batteries.

***Genetic Analysis***

For the complete list of faecal samples used in this study please see Table A1.

*DNA extraction*

DNA was extracted using the QIAamp Fast DNA Stool Mini Kit (Qiagen) following the human DNA analysis protocol, with minor procedures modifications: (i) ~250 uL of the faecal slurry were added to 1 ml InhibitEX buffer, shaking for 5 min at 20 Hz of speed ug Tissue Lyser II (Qiagen), before proceeding to step 3; (ii) in step 14 100 uL of Buffer ATE added instead of 200 uL to increase DNA yield.

*Mitochondrial DNA*

Amplification of the mitochondrial control region (460 bp) was conducted using the species-specific Mono1 (forward primer 5’-CTACTCCCTGAATAACCAAC-3’) and Mono2 (reverse primer 5’-AGCGAGAAGAGCGGCAAATG-3’) (Fagundes et al., 2008). PCR reactions were carried out in a total volume of 20 uL containing 2 uL of DNA extract, 0.5 uL of each primer, and 17 uL of MyTaq Red Mix (Bioline). Amplifications were performed at 95°C for 2 min, followed by 35 cycles of 92°C for 60 s, 48°C for 30 s and 72°C for 45 s, with a final extension at 72°C for 5 min. The PCR products were quantified using 1.5% of TBE agarose gels with HyperLadder 50bp (Bioline). Standard Sanger sequencing was conducted bidirectionally by Macrogen Inc., Korea. Obtained sequences were checked and assembled in DNA Baser Assembler v5.15 (2014, Heracle BioSoft, www.DnaBaser.com). Molecular species identification was conducted through a BLAST search in the NCBI database. Then, sequences were aligned using the CLUSTAL W and visually inspected in MEGA version X (Kumar et al. 2018), using a *Brachyteles hypoxanthus* reference sequence retrieved from GenBank (AF213966).

*Microsatellite genotyping*

For the microsatellite analyses, a set of 14 loci were amplified through a combination of two or three primers labelled with fluorescent dyes in multiplexed PCR reactions (Table A2). PCR reactions were performed in a final volume of 20 uL, containing 2 uL of DNA extract, 0.4 uL of each primer (10 uM), 10 uL of Multiplex PCR Master Mix (Qiagen), and H_2_O. Negative PCR blanks (i.e., without DNA template) were used as a control for each reaction. Thermal cycling conditions were as follows: initial denaturation at 95°C for 15 min, followed by 34 cycles of 94°C for 30s, 55°C for 90s and 72°C for 60s, with a final extension at 60°C for 30 min. Automated capillary analyses were conducted on an ABI 3730XL DNA Analyser using GeneScan 400HD size standard marker (Applied Biosystems) for allele size estimates (Macrogen Inc., Korea).

All analysed samples were genotyped at least three times for all loci, to minimise genotyping errors.  Automated allele calling was conducted in GeneMapper v.4.1 (Applied Biosystems) and further visually checked for genotype determination. Genotypes were independently determined twice for all samples per replicate (Stojak et al. 2016). Homozygote genotypes were considered when scored on all replicates or in at least two out of the three replicates when one replicate amplification failed. Heterozygote was accepted if scored on two out of three replicates. If none of these criteria were met, alleles were classified as missing data.

The probability of identify (PI) measures for unrelated (PI*_ave_*) and siblings (PI*_sibs_*) were calculated in GenAlex (Peakall & Smouse, 2006). Genetic tagging of individuals was as follow:

1) Distinct individuals:

a) possessing different mtDNA haplotype and same/distinct microsatellite genotypes,
b) possessing mtDNA haplotype and differing microsatellite genotypes.

2) Same individual:

a) whether presenting matches for both genotypes and mtDNA haplotypes. All samples identified with the same haplotype and a matching score ≥87.5% loci were re-examined for possible genotyping errors.

Since we sampled unhabituated muriquis and the genetic sampling was conducted in distinct sampling sessions, individuals could be sampled “recaptured” several times. We considered independence of genetic sampling events as follows (Arandjelovic et al. 2010; McCarthy et al. 2015):

a) distinct samples from the same individual were considered independent if were separately geographically and/or temporally in distinct days;
b) distinct samples from the same individual were considered duplicate if they were collected in the same day and a distance apart of <3479 m (based on the maximum travel distance of muriquis within one day, Lima et al. 2019).

Table A 1. Northern muriqui samples information, including sample ID, type of sample, date of collection and DNA extraction, geographical coordinates and respective haplotype inferred by mtDNA markers, and genotypes matches (Duplicate: same individual collected twice during the same day; excluded: not included in the microsatellite analysis).

| Sample ID | Group | Sample Type | Lat | Long | Collection Date | Extraction Date | Hap | Genotype matches |
| --- | --- | --- | --- | --- | --- | --- | --- | --- |
| PNC02 | VA | Fecal | -20.47969 | -41.84179 | 23-Jan-17 | Nov-18 | h14 |  |
| PNC03 | VA | Fecal | -20.47969 | -41.84179 | 23-Jan-17 | Nov-18 | h14 | duplicate (PNC02) |
| PNC04 | VA | Fecal | -20.47969 | -41.84179 | 23-Jan-17 | Nov-18 | h14 |  |
| PNC05 | VA | Fecal | -20.47969 | -41.84179 | 23-Jan-17 | Nov-18 | h14 |  |
| PNC06 | VA | Fecal | -20.47969 | -41.84179 | 23-Jan-17 | Nov-18 | h14 |  |
| PNC07 | VA | Fecal | -20.47969 | -41.84179 | 23-Jan-17 | Nov-18 | h14 |  |
| PNC08 | VA | Fecal | -20.47969 | -41.84179 | 23-Jan-17 | Nov-18 | h19 |  |
| PNC09 | VA | Fecal | -20.47969 | -41.84179 | 23-Jan-17 | Nov-18 | h19 |  |
| PNC10 | VA | Fecal | -20.47969 | -41.84179 | 23-Jan-17 | Nov-18 | h14 |  |
| PNC11 | VA | Fecal | -20.48025 | -41.84542 | 06-Fev-17 | Nov-18 | h19 |  |
| PNC12 | VA | Fecal | -20.48164 | -41.85045 | 07-Feb-17 | Nov-18 | h19 |  |
| PNC13 | VA | Fecal | -20.48164 | -41.85045 | 07-Feb-17 | Nov-18 | h1 | excluded |
| PNC14 | VA | Fecal | -20.48164 | -41.85045 | 12-Feb-17 | Nov-18 | h8 |  |
| PNC17 | VSM | Fecal | -41.74243 | -20.49017 | 14-Fev-17 | Nov-18 | h9 |  |
| PNC18 | VSM | Fecal | -41.74904 | -20.49035 | 17-Fev-17 | Nov-18 | ns |  |
| PNC21 | VSM | Fecal | -41.75539 | -20.49147 | 21-Fev-17 | Nov-18 | h1 |  |
| PNC22 | VSM | Fecal | -41.75539 | -20.49147 | 21-Fev-17 | Nov-18 | h9 |  |
| PNC24 | VSM | Fecal | -41.75539 | -20.49147 | 21-Fev-17 | Feb-19 | h1 |  |
| PNC25 | VSM | Fecal | -41.75539 | -20.49147 | 21-Fev-17 | Nov-18 | h1 |  |
| PNC28 | VA | Fecal | -41.84696 | -20.48062 | 15-Mar-17 | Nov-18 | h1 |  |
| PNC29 | VSM | Fecal | -41.73706 | -20.49174 | 17-Mar-17 | Nov-18 | h11 |  |
| PNC30 | VSM | Fecal | -41.73706 | -20.49174 | 17-Mar-17 | Nov-18 | h14 |  |
| PNC31 | VSM | Fecal | -41.73935 | -20.49052 | 23-Mar-17 | Nov-18 | h9 |  |
| PNC32 | VSM | Fecal | -41.73935 | -20.49052 | 23-Mar-17 | Nov-18 | h11 |  |
| PNC33 | VSM | Fecal | -41.73935 | -20.49052 | 23-Mar-17 | Nov-18 | h9 |  |
| PNC34 | VSM | Fecal | -41.73935 | -20.49052 | 23-Mar-17 | Nov-18 | h9 | duplicate (PNC31) |
| PNC35 | VSM | Fecal | -41.75174 | -20.48993 | 17-Apr-17 | Nov-18 | h14 |  |
| PNC38 | VA | Fecal | -41.84536 | -20.48004 | 09-May-17 | Nov-18 | h1 |  |
| PNC39 | VA | Fecal | -41.84556 | -20.48000 | 09-May-17 | Nov-18 | h19 |  |
| PNC41 | VSM | Fecal | -41.75754 | -20.49206 | 10-Jun-17 | Nov-18 | h2 |  |
| PNC75 | VA | Fecal | -41.85667 | -20.48472 | 20-Sep-17 | Nov-18 | h17 |  |
| PNC76 | VA | Fecal | -41.85667 | -20.48472 | 20-Sep-17 | Nov-18 | h14 |  |
| PNC78 | VA | Fecal | -41.85444 | -20.48222 | 16-Fev-18 | Nov-18 | h9 |  |
| PNC79 | VA | Fecal | -41.84944 | -20.48111 | 17-Fev-18 | Nov-18 | h14 |  |
| PNC80 | VA | Fecal | -41.84944 | -20.48111 | 17-Fev-18 | Nov-18 | h1 |  |
| PNC81 | VA | Fecal | -41.84944 | -20.48111 | 17-Fev-18 | Nov-18 | h1 |  |
| PNC82 | VA | Fecal | -41.84944 | -20.48111 | 17-Fev-18 | Nov-18 | h14 |  |
| PNC83 | VA | Fecal | -41.84944 | -20.48111 | 17-Fev-18 | Nov-18 | h14 | excluded |
| PNC84 | VA | Fecal | -41.84944 | -20.48111 | 17-Fev-18 | Nov-18 | h1 |  |
| PNC85 | VA | Fecal | -41.84944 | -20.48111 | 17-Fev-18 | Nov-18 | h14 |  |
| PNC86 | VA | Fecal | -41.84944 | -20.48111 | 17-Fev-18 | Nov-18 | h14 |  |
| PNC87 | VA | Fecal | -41.84944 | -20.48111 | 17-Fev-18 | Nov-18 | h1 |  |
| PNC88 | VA | Fecal | -41.84889 | -20.48139 | 17-Fev-18 | Nov-18 | h14 | excluded |
| PNC89 | VA | Fecal | -41.84944 | -20.48111 | 17-Fev-18 | Nov-18 | h14 |  |
| PNC90 | VA | Fecal | -41.84944 | -20.48111 | 17-Fev-18 | Nov-18 | h14 |  |
| PNC91 | VA | Fecal | -41.84944 | -20.48111 | 17-Fev-18 | Nov-18 | h14 |  |
| PNC92 | VA | Fecal | -41.84944 | -20.48111 | 17-Fev-18 | Nov-18 | h20 |  |
| PNC93 | VA | Fecal | -41.84944 | -20.48111 | 17-Fev-18 | Nov-18 | h14 |  |
| PNC102 | VA | Fecal | -41.75618 | -20.48358 | 07-Mar-18 | Nov-18 | h14 |  |
| PNC103 | VA | Fecal | -41.75699 | -20.48369 | 07-Mar-18 | Nov-18 | h19 |  |
| PNC104 | VA | Fecal | -41.84822 | -20.48088 | 07-Mar-18 | Nov-18 | h15 |  |
| PNC106 | VA | Fecal | -41.85196 | -20.48203 | 07-Mar-18 | Nov-18 | h19 |  |
| PNC107 | VA | Fecal | -41.84300 | -20.48764 | 12-Mar-18 | Nov-18 | h20 |  |
| PNC108 | VA | Fecal | -41.84346 | -20.48770 | 12-Mar-18 | Nov-18 | h20 |  |
| PNC109 | VA | Fecal | -41.84293 | -20.48722 | 12-Mar-18 | Nov-18 | h21 |  |
| PNC110 | VA | Fecal | -41.84293 | -20.48722 | 12-Mar-18 | Nov-18 | h21 |  |
| PNC111 | VA | Fecal | -41.84293 | -20.48722 | 12-Mar-18 | Nov-18 | h7 |  |
| PNC112 | VA | Fecal | -41.84293 | -20.48722 | 12-Mar-18 | Feb-19 | ns |  |
| PNC113 | VA | Fecal | -41.75618 | -20.48358 | 05-May-18 | Feb-19 | h7 |  |
| PNC114 | VA | Fecal | -41.75618 | -20.48358 | 05-May-18 | Feb-19 | h8 |  |

Table A 2. Primers set and description of microsatellites loci studied for northern muriqui (Brachyteles hypoxanthus) population at Caparaó National Park, Brazil, including repeat motif, primer sequence, fluorescent dye, number of PCR replicate, size-range in this study and in the literature, species origin and primers reference.

| **Locus** | **Repeat Motif** | **Primers (5’→3’)** | **Dye** | **Replicate** | **Size-range this study** | **Size-range*** | **Species origin** | **Reference** |
| --- | --- | --- | --- | --- | --- | --- | --- | --- |
| Multiplex I | | |  |  |  |  |  |  |
| LL113 | (GT)_15_ | GCAAAACTCCCCTGTGACTG | FAM | 4 | 167-189 | 181–186 | *Lagothrix lagotricha* | 1 |
|  |  | CCCACTCTCCTCCACAAAGG |  |  |  |  |  |  |
| SB30 | (CA)_9_-AT-(CA)_11_ | TAAAGTTAAGATTGGATTTCAC | FAM | 4 | 101-111 | 83-103 | *Saguinus bicolor* | 2 |
|  |  | GCAGAAAAACCTAACAATACA |  |  |  |  |  |  |
| D5S111 | CA | GGCATCATTTTAGAAGGAAAT | HEX | 5 | 161-171 | 162-174 | *Homo sapiens* | 3 |
|  |  | ACATTTGTTCAGGACCAAAG |  |  |  |  |  |  |
| Multiplex II | | |  |  |  |  |  |  |
| LL1110 | (GT)_2_ | GGTGAATGAGAGAATCAAAG | HEX | 4 | 212-218 | 218-222 | *Lagothrix lagotricha* | 1 |
|  |  | TATGTTCCACAGTAGAAAGC |  |  |  |  |  |  |
| SB38 | (CA)_19_ | GCCTCAATGGGTTTTAACC | FAM | 3 | 125-139 | 126-144 | *Saguinus bicolor* | 2 |
|  |  | AGAACGAGTCTGTATCTTGA |  |  |  |  |  |  |
| D17S804 |  | GCCTGTGCTGCTGATAACC | HEX | 5 | 151-159 | 152-162 | *Homo sapiens* | 4 |
|  |  | CACTGTGATGAGATGTCATTCC |  |  |  |  |  |  |
| Multiplex III | | |  |  |  |  |  |  |
| LL1115 | (GT)_3_(GA)_1_(GT)_5_ | GCTCATATTCATACATCCCTTGG | HEX | 4 | 194-206 | 196–208 | *Lagothrix lagotricha* | 1 |
|  |  | TTTGCTTGCTCATTCATTGC |  |  |  |  |  |  |
| LL1118 | (CA)_2_(TA)_1_(CA)_17_ | TTTCTCCCTCTCAGATTACCAG | FAM | 6 | 128-156 | 142–166 | *Lagothrix lagotricha* | 1 |
|  |  | CCTTGAGGTTTTTGGGTTCC |  |  |  |  |  |  |
| Multiplex IV | | |  |  |  |  |  |  |
| D8S165 | (CA)_18_ | ACAAGAGCACATTTAGTCAG | TET | 4 | 126-148 | 123-146 | *Homo sapiens* | 5 |
|  |  | AGCTTCATTTTTCCCTCTAG |  |  |  |  |  |  |
| LL157 | (GA)_4_(GT)_4_(CT)_1_ | TGGCAAGTCTGGTTTCAAGC | TET | 3 | 223-233 | 224-232 | *Lagothrix lagotricha* | 1 |
|  |  | TTCCAGACTGAGCTAGGATGC |  |  |  |  |  |  |
| LEON15 | (GA)_17_ | CTGATCCTTGAAGCAGCATTG | TET | 3 | 257-269 | 259-269 | *Leontopithecus chrysopygus* | 6 |
|  |  | GGTTAAAGGGGTTCGTTCTGTG |  |  |  |  |  |  |
| Multiplex V | | |  |  |  |  |  |  |
| LrP2BH6 | (CA)_19_ | TCTGTTTGAATCCCCAGTCC | FAM | 4 | 96-116 | 92-114 | *Leontopithecus rosalia* | 7 |
|  |  | GCAGTCCCTCAAGGTTTTCT |  |  |  |  |  |  |
| AB06 | (CT)_9_TTT(CT)_11_GTCTGTCTTAT(AC)_16_ | GTGATTATTGTGTGGTACTTG | TET | 3 | 257-279 | 261-273 | *Alouatta belzebul* | 8 |
|  |  | ATGTATTTTTCTGGTTTTACC |  |  |  |  |  |  |
| Multiplex VI | | |  |  |  |  |  |  |
| LEON21 | (GT)_19_(NA)_1_(GT)_5_ | CAGTTGAGGGAACAGGAATTA | FAM | 4 | 365-383 | 367-379 | *Leontopithecus chrysopygus* | 6 |
|  |  | CACTGCACTGACAGAGCAAG |  |  |  |  |  |  |
| *Strier, K. B., Chaves, P. B., Mendes, S. L., Fagundes, V., & Di, A. (2011). Low paternity skew and the influence of maternal kin in an egalitarian, patrilocal primate. PNAS 108 (47), 18915-18919.  1. Di Fiore A, Fleischer RC (2004) Microsatellite markers for woolly monkeys (Lagothrix lagotricha) and their amplification in other New World primates (Primates: Platyrrhini). Mol Ecol Notes 4:246e249.  2. Bohle UR, Zischler H (2002) Polymorphic microsatellite loci for the mustached tamarin (Saguinus mystax) and their cross-species amplification in other New World monkeys. Mol Ecol Notes 2:1e3.  3. Weber JL, Kwitek AE, May PE (1990) Dinucleotide repeat polymorphisms at the D5S107, D5S108, D5S111, D5S117 and D5S118 loci. Nucleic Acids Res 18:4035.  4. Weissenbach J, et al. (1992) A second-generation linkage map of the human genome. Nature 359:794e801. | | | | | | | | |
| 5. Weber JL, May PE (1989) Abundant class of human DNA polymorphisms which can be typed using the polymerase chain reaction. Am J Hum Genet 44:388e396 | | | | | | | | |
| 6. Perez-Sweeney BM, et al. (2005) Dinucleotide microsatellite primers designed for a critically endangered primate, the black lion tamarin (Leontopithecus chrysopygus). Mol Ecol Notes 5:198e201. | | | | | | | | |
| 7. Grativol AD, Ballou JD, Fleischer RC (2001) Microsatellite variation within and among recently fragmented populations of the golden lion tamarin (Leontopithecus rosalia). Conserv Genet 2:1e9. | | | | | | | | |
| 8. Goncalves EC, Silva A, Barbosa MSR, Schneider MPC (2004) Isolation and characterization of microsatellite loci in Amazonian red-handed howlers Alouatta belzebul (Primates, Plathyrrini). Mol Ecol Notes 4:406e408 | | | | | | | | |
